# Algae–bacteria associations provide metabolite-mediated protection against algicidal bacteria in a tripartite plankton community

**DOI:** 10.64898/2026.08.28.747787

**Authors:** Shahrukh A. Siddiqui, Christian Zerfaß, Vera Nikitashina, Ruyi Yu, Georg Pohnert

## Abstract

Microalgal fitness in nature is shaped by interactions within a diverse microbial community, yet most experimental studies have examined algal-bacterial interactions in pairwise systems. It is well established that bacteria can exhibit growth promoting or inhibiting effects on co-existing algae. Comparatively little information is available about how additional partners can alter the outcome of diatom-bacteria interactions. In the present study, we screened the pairwise interaction of the marine diatom *Skeletonema marinoi* with ten different bacteria. This screening identified *Marinobacter adhaerens* as a growth promoting and *Vibrio cyclitrophicus* HSW24 as growth inhibiting partner. Growth inhibition of *V. cyclitrophicus* was associated with cell lysis, chain fragmentation and altered pigmentation whereas *M. adhaerens* supported increased chlorophyll *a* fluorescence, uniform pigmentation, intact chains and healthy cell morphology. In a tripartite community containing both bacteria and the alga, *M. adhaerens* protected *S. marinoi* from the inhibitory effect of *V. cyclitrophicus* in a density dependent manner. Comparative metabolomics revealed distinct metabolic profiles between the pairwise and tripartite interactions. This allowed to identify metabolites that were up-regulated in the tripartite community and therefore candidates for the observed protection. Among these, kynurenic acid and *N*-acetyltyramine were identified in bioassays as protective molecules, thus clearly highlighting the importance of secondary metabolites in this interaction. The present findings demonstrate that a third bacterial partner can alter the outcome of an antagonistic algal-bacterial interaction by means of chemical mediators. This work has implications for our understanding of microbial community functioning that cannot only be derived from the investigation of pairwise interactions.

## INTRODUCTION

Diatoms are unicellular photosynthetic microorganisms and form the foundation of the marine food web. They are responsible for approximately 20-25% of global primary production thereby playing an important role in the global carbon cycle [1, 2]. Diatoms release substantial amounts of dissolved organic matter into the seawater which is utilized by heterotrophic bacteria. These bacteria can influence diatom growth, metabolism, physiology and ecological functioning and thus influence development and demise of diatom blooms [3–5]. Many of these interactions occur in the phycosphere, a microenvironment surrounding diatoms which is rich in organic matter and characterized by steep gradients of nutrients that promote bacterial chemotaxis, attachment and metabolic cross feeding [6]. Diatoms actively shape the phycosphere by releasing metabolites that can serve as substrate, antimicrobial factors, or chemical mediators that control bacterial function and abundance [7, 8].

The role of bacteria in modulating diatom growth is well established, although research has predominantly focused on pairwise interactions. These studies have demonstrated that beneficial bacteria can enhance algal growth through various mechanisms, including vitamin provision, nutrient remineralization, iron mobilization, and the production of hormone-like compounds. For example, bacterial partners such as Sulfitobacter provide growth promoting indoleacetic acid to *Pseudo-nitzschia, Ruegeria* sp. can provide the essential vitamin-B12 to *Thalassiosira pseudonana* and *Marinobacter* sp. can supply iron to diatoms. Bacterial diketopiperazine peptides promote the growth of *Phaeodactylum tricornutum* [6, 8, 9]. On the other hand, harmful or pathogenic bacteria can inhibit algal growth or induce cell death [7, 10, 11]. Such algicidal bacteria can act through direct contact or secretion of harmful factors including lytic compounds, disruptors of photosynthesis or causative agents of oxidative stress. These can result in algal bloom termination thereby shaping phytoplankton succession [4, 12, 13]. The mode of action of algicidal bacteria is diverse. For example, *Kordia algicida* lyses the diatom *Skeletonema marinoi* by means of a secreted protease AlpA in combination with low-molecular-weight metabolites; *Vibrio coralliirubri* employs not fully characterized contact dependent mechanisms and kills *Karenia mikimotoi* whereas *Pseudomonas* sp. uses indolinedione derivatives to kill *Chaetoceros ceratosporum* [11, 14, 15].

Pairwise interaction studies have revealed important mechanisms, but they are not sufficient to explain the ecological complexity of marine microbial communities. In nature, diatoms rarely encounter a single bacterial strain in isolation rather they interact with multiple microbial partners simultaneously in the phycosphere [3]. These communities are stabilized by exchange of nutrients, signalling molecules and inhibitory metabolites which suggests community level outcomes that cannot always be predicted from pairwise interactions alone [3, 11]. Despite increasing recognition that microbial interactions are shaped by community context, it remains unclear whether a beneficial bacterial partner can suppress or redirect algicidal activity in a marine diatom, and which chemical processes mediate such protection.

Evidence from freshwater algae suggest that higher order interactions can protect hosts from pathogenic bacteria. For example, introduction of third microbial partner *Aspergillus nidulans* protects *Chlamydomonas reinhardtii* from algicidal activity of *Streptomyces iranensis* [16]. *Chlamydomonas* is also protected from algicidal lipopeptide produced by *Pseudomonas protogens* by the bacterium *Mycetocola lacteus* [17].

In marine diatoms the frequency and mechanisms of such multipartite interactions are not known. Therefore, we tested here the hypotheses that an additional bacterial partner can alter the outcome of pairwise algae-bacteria interactions and that specialized metabolites are mediating the involved processes. We used a bloom forming diatom *S. marinoi*, a well-established model species, which is widely distributed in temperate coastal waters [18–20]. Based on an initial screening, we selected *Marinobacter adhaerens* as a growth promoting and *Vibrio cyclitrophicus*-HSW24 as a growth inhibitory bacterium. Our findings highlight the complexity of diatom-bacteria interactions and the important roles of specialized metabolites in marine microbial communities.

## MATERIAL AND METHODS

### Solvents Used

All solvents used in the present study for endo- and exo-metabolome extraction, liquid chromatography high resolution mass spectrometry and standard preparation were of mass spectrometry (MS) grade. The solvents used in the present study comprised water (Chromasolv Plus for HPLC, Honeywell, Germany), methanol (SupraSolv, Merck, Germany), chloroform (HPLC grade, FisherScientific, UK), ethanol (LiChrosolv, Supelco, Merck, Germany), and acetonitrile (CHEMSOLUTE, Th. Geyer, Germany).

### Cultivation of Skeletonema marinoi

*S. marinoi* (RCC75) was procured from the Roscoff culture collection (Roscoff, France) and cultivated in Artificial Sea Water (ASW) [21]. To reduce the abundance of associated bacteria, cultures were treated with antibiotics following [22] with minor modifications. *S. marinoi* cells were collected on a 5 µm cell strainer (PluriSelect) and washed with 10 mL of Triton X-100 solution (20 µg/mL) to remove attached bacteria. The cells were washed at least twice with ASW to remove residual Triton X-100. The cultures were then treated with antibiotic cocktails (Kanamycin 50 µg mL^-1^, Ciprofloxacin 20 µg mL^-1^ and Chloramphenicol 2 µg mL^-1^) for 24 hours. Following treatment, cells were recovered by centrifugation at 3,500 g for 5 min and washed with ASW to remove remaining antibiotics. The cells were suspended in fresh ASW and incubated for 8-10 days at 13 °C under diurnal light : dark cycle of 14 : 10h with a light intensity of 25 µmol photon m^-2^ s^-1^. To assess the presence of culturable bacteria, 50 µL of treated cultures were inoculated directly into marine broth (MB) and an additional 50 µL were spread on marine agar (MA) plates and incubated at 28 °C for 48 h-72 h. Bacterial growth was assessed by monitoring turbidity in the MB or visible bacterial colonies on MA plates. Cultures showing no observed bacterial growth were maintained under previously described culture conditions and used in further experiments. The absence of bacteria was not verified by staining or sequencing therefore, the cultures were considered to have reduced bacterial load rather than to be axenic.

### Bacteria isolation and identification

In the present study, we screened a total of 10 widely distributed bacterial strains (Supporting Table 1) for their effects on the growth of *S. marinoi.* Out of these, seven strains had previously been maintained in our laboratory, two were obtained from DSMZ culture collection (Germany) whereas, one was isolated from seawater from Helgoland, Germany.

The bacterial isolation was done by serial dilution of Helgoland plankton samples and plating it on MA plates. The plates were incubated at 28 °C for 7 days. Morphologically distinct single colonies were picked and repeatedly streaked to fresh MA plates three to four times to obtain a pure bacterial culture. A single colony was transferred to MB and incubated at 28 °C with continuous shaking at 120 rpm. Once the OD_600_ reached ∼0.6, the cultures were harvested and genomic DNA was extracted using a bacterial DNA isolation kit (Jena Biosciences) following the manufacturer’s protocol.

The 16s rRNA gene was amplified through polymerase chain reaction (PCR) from genomic DNA using universal primer pair 27F (5′-AGAGTTTGATCCTGGCTCAG-3′) and 1492R (5′-ACGGHTACCTTGTTACGACTT-3′). The PCR reaction mixture was prepared in 25 µL and contained 0.5 µM each primer, 2.5 µL of 10x reaction buffer, 250 µM dNTPs, 1.25 U Taq polymerase, 25 ng of template (DNA) and the final volume was reconstituted with ultrapure autoclaved water. The PCR reaction condition constituted of initial denaturation at 95 °C for 5 min followed by the final denaturation at 95 °C for 30 sec, annealing at 56 °C for 30 sec and extension at 72 °C for 60 sec for 30 cycles with an additional final extension at 72 °C for 10 min. The PCR product was verified on 1% agarose gel and was purified using a PCR purification kit (Qiagen) following the manufacturer’s protocol. The purified PCR product was sent to Eurofins, (Germany) for Sanger sequencing. The resulting 16S rRNA gene sequence was submitted to the NCBI nucleotide database using BLASTn for taxonomic identification. A Phylogenetic tree was constructed based on maximum likelihood analysis using MEGA12.0 software.

### Interaction of *S. marinoi* with bacteria

To screen for growth promoting and growth inhibiting bacteria cultures were prepared by inoculating a single colony from a MA plate into 3 mL MB and incubation at 28 °C with continuous shaking at 120 rpm. Once the OD_600_ reached ∼0.6, bacteria were transferred into fresh MB at a 1:100 v/v dilution and incubated overnight under the same conditions. When the OD_600_ reached 0.5-0.7, the bacterial cultures were transferred to 1.5 mL microcentrifuge tubes (Eppendorf) and centrifuged at 10,000 rpm for 2 minutes at room temperature. Bacterial cell pellets were washed twice with ASW to completely remove MB and then resuspended in fresh ASW.

For pairwise co-cultivation, *S. marinoi* from mid exponential phase were inoculated in ASW in tissue culture flasks and the initial chlorophyll *a* fluorescence was adjusted to 0.1. A bacterial inoculum was added to give the final concentration of 1 x 10^5^ - 10^6^ CFU mL^-1^. The co-cultures were incubated at 13 °C under a 14 : 10 h light : dark cycle. Simultaneously, a process blank consisting of only ASW was incubated under identical conditions. Chlorophyll *a* fluorescence was used as a proxy for *S. marinoi* growth and was recorded every second day using a Varioskan Flash Multimode Reader (Thermo Scientific) by transferring 200 uL of culture to a 96 well plate. Microscopic examination was also done every second day using an inverted microscope (Leica DM LED; Leica Microsystems, Wetzlar, Germany) to study cell morphology.

### Tripartite synthetic community interaction

Aliquots of exponentially growing *S. marinoi* cultures (9-10 days old) were transferred to fresh TC flasks and the initial chlorophyll *a* fluorescence was adjusted to 0.1 with ASW. Fresh bacterial cultures of *V.* cyclitrophicus and *M. adhaerens* were prepared as described above. Once the bacterial OD_600_ was ∼0.6, the cells were harvested, washed to remove MB and added to *S. marinoi* cultures to give the final concentration of approximately 10^3^-10^4^ CFU mL^-1^. The cultures were incubated as described above. We established tripartite communities using combinations of *M. adhaerens* and *V. cyclitrophicus* for inoculation in ratios of 1:1, 2:1, and 3:1. The chlorophyll *a* fluorescence was measured every second day as a proxy for *S. marinoi* growth.

### Sample preparation for non-targeted metabolomics

*S. marinoi* cells were microscopically counted on day 10 and equal number of cells from all samples were transferred to 50 mL falcon tubes and centrifuged at 2,500 rpm for 15 min. The supernatant was collected in fresh falcon tubes and used for exo-metabolome extraction whereas the cells pellet was washed with fresh ASW and used for endo-metabolite extraction.

Endo-metabolites were extracted following [23] with some modifications. Cell pellets were resuspended in 1 ml of an ice cold extraction solvent (methanol: ethanol: chloroform; 1:3:1; v:v:v). Samples were vortexed for 1 min and sonicated at 25 °C for 15 min. The extracts were centrifuged at 13,000 rpm for 15 min. The resulting supernatant was collected, dried under nitrogen stream and stored at -20 °C until analysis.

For exo-metabolome extraction, the supernatant of the above centrifugation was loaded on an Oasis PRiME HLB solid phase extraction cartridge (3 cc, Waters) and processed according to manufacturer’s instruction. After elution, the methanolic extracts were dried under nitrogen a stream and stored at -20 °C. Blanks consisting of only ASW were processed in the same manner alongside the samples.

Before LC-MS analysis, the dried extracts were resolubilized in 200 µL of methanol and centrifuged again at 13,000 rpm for 15 min. Pooled quality control (QC) sample was prepared by mixing 10 µl from all the samples except process blanks.

### LC-MS/MS Analysis

Extracts were analyzed using a Vanquish LC system equipped with autosampler and coupled to an Orbitrap Exploris 480 mass spectrometer (Thermo Fisher Scientific). The full MS1 scans were acquired in polarity switching mode whereas MS/MS spectra were acquired in positive ion mode using the AcquireX (AcqX) workflow.

One µl of each sample was injected and separated using an Accucore C18 column (100 × 2.1 mm, 2.6 µm, Thermo Fisher Scientific, Bremen, Germany) with a flow rate of 0.4 mL min^-1^. Mobile phase A consisted of water containing 2% acetonitrile and 0.1% formic acid whereas, mobile phase B consisted of 100% acetonitrile. The gradient started with 100 % A for 0.2 min followed by an increase to 100% B within 8 min. This was held for 3 min before returning to 100 % A (2 min) at a flow of 0.4 mL min^-1^. The column oven temperature was 25°C. Full MS1 spectra were acquired with a scan range of *m*/*z* 80 to 1200 at a resolution of 70,000 using electrospray ionization. For MS/MS acquisition, an inclusion list was created using the AcquireX workflow by running pool-QC sample. Five injections were performed to increase MS/MS coverage. Precursor ions were selected with an isolation window of 0.4 *m*/*z* and the AGC target was set to 3 × 10^6^.

Raw data were processed through Compound Discoverer version 3.3 (Thermo Fischer Scientific) with the minimum peak quality rating of 6 detected in a minimum of 50% of samples. Peak picking, peak deconvolution, chromatographic alignment, background subtraction and metabolite annotation were performed with the software. Precursor mass tolerance was set to 5 ppm, minimum peak intensity threshold to 10,000 and the average peak area of sample was set to at least 5 fold higher than that in blank samples. The masses were searched using compound discoverer 3.3 (LIPID MAPS, Natural Product Atlas and an in house library) and MS/MS spectra were additionally mapped against the mzCloud spectral library. Raw data were also converted to mzXML format using ProteoWizard and MS/MS spectra were also analyzed using SIRIUS. Metabolites annotated by spectral library matching were considered putatively identified unless mentioned as confirmed using authentic standards.

### Data and Statistical analysis

All experiments were performed with a minimum of four biological replicates. The feature table generated in Compound Discoverer was processed and analyzed using MetaboAnalyst software version 6.0 for statistical analysis. Data were log-transformed and pareto-scaled prior to multivariate analysis. PCA was performed to evaluate the variation among treatments. Differentially abundant features were visualized using volcano plots, with significant features defined by a fold change greater than 2 and false discovery rate adjusted *p*-value <0.05.

## RESULTS

### Bacterial isolation, phylogeny and effect on *S. marinoi* growth

We selected 9 widely distributed marine bacteria and a *Virbrio* species that was isolated during a diatom bloom in Helgoland, co-occurring with *S. marinoi*. Sequence similarity analysis of 16S rRNA using NCBI nucleotide database (BLAST) identified the isolated bacterial strain as *Vibrio cyclitrophicus* HSW24 with 100% query coverage and 99.6% sequence identity to the closest database match. The sequence has been submitted in the NCBI GenBank database with accession number PZ671326.

Phylogenetic analysis of the 16S rRNA gene sequences indicated that the bacterial isolates were phylogenetically diverse and grouped broadly according to their respective taxonomic lineages (Supporting Fig. 1). *M. adhaerens* clustered with other *Marinobacter* strains, whereas *V. cyclitrophicus* formed a separate branch. The two *Phaeobacter* strains clustered closely together and the remaining isolates formed separate genus level lineages. This analysis suggests that the bacterial screening covered a taxonomically diverse set of marine bacteria.

**Figure 1.**
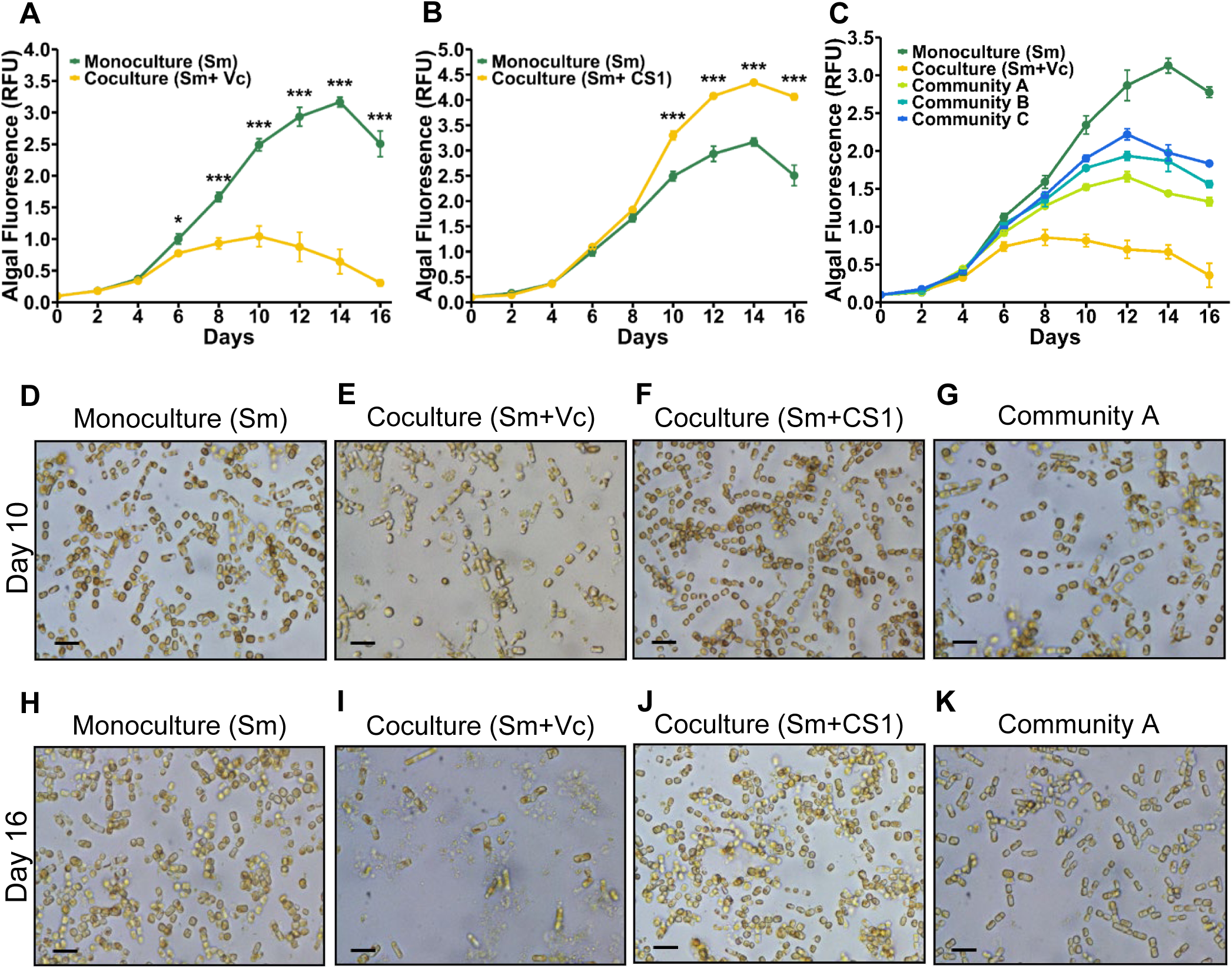
Modulation of *S. marinoi* (Sm) growth by bacteria in coculture and community. (A-C) Fluorescence based growth curve of Sm in monoculture, coculture with Vc (A), coculture with CS1 (B) and in a community consist of Sm, CS1 and Vc (C). In the community treatments, the initial Sm cell density was kept constant, while the abundance of CS1 was increased relative to Vc. Community A contained Vc:CS1 at a 1:1 cell number ratio, Community B at 1:2, and Community C at 1:3. Data are presented as mean ± SE from 5 biological replicates. Differences at individual time point are assessed using two-tailed Student’s t-test; *\*p<0.05, ***p<0.001*. (d-k) Representative microscopy images of Sm in monoculture (D,H), coculture with Vc (E,I), coculture with CS1 (F, J) and in the community A (G,K). Scale bars represent 20 µm. The full set of microscopy images from day 2 to day 16 is provided in Supporting Fig. 3. **Sm**, *Skeletonema marinoi*; **Vc**, *Vibrio cyclitrophicus* HSW24; **CS1**, *Marinobacter adhaerens* CS1.

The effect of the respective bacteria on *S. marinoi* growth was monitored in a pairwise coculture using Chlorophyll *a* fluorescence as a proxy for algal growth (Supporting Fig. 2 A-J). *Croceibacter* sp. (Cro1) caused significant growth inhibition of *S. marinoi* from day 4 onwards whereas, *V. cyclitrophicus* inhibited *S. marinoi* growth from day 6. *Pseudoalteromonas* sp. (CR1) initially supported *S. marinoi* growth but from day 10 onward it also caused significant growth inhibition. *Phaeobacter inhibens* (Pha) and *M. adhaerens* significantly promoted *S. marinoi* growth, with the effect being more pronounced for *M. adhaerens*. *Phaeobacter glacielencis* (Pg), *Roseovarious* sp. and *Maribacter* (Mari1) also slightly supported growth. *Marinobacter sp*. (Marino1) showed a mixed effect i.e., promoting growth in early growth phase and growth inhibition during late growth phase whereas, *Marinobacter algicola* (Ma) showed no effect on *S. marinoi* growth (Supporting Fig. 2).

**Figure 2.**
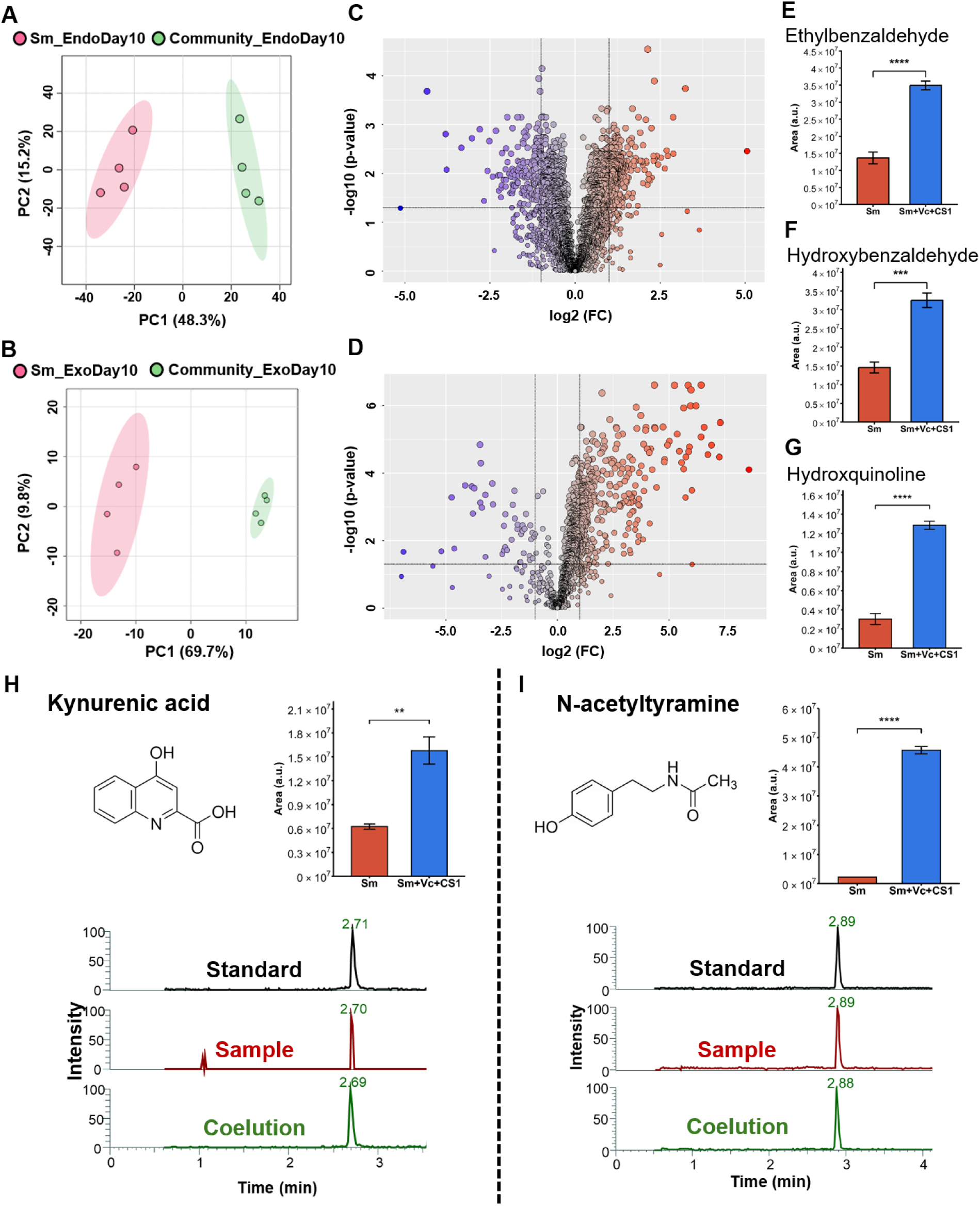
Metabolic changes of the protective tripartite community. (A,B) Principal Component Analysis score plot of endo-metabolome (A) and exo-metabolome (B) in Sm (control) and community (tripartite community; Sm+Vc+CS1). (C,D) Volcano plots showing differentially regulated features in the endo-metabolome (C) and exo-metabolome (D). (E-G) Relative peak areas of selected differentially abundant features putatively annotated based on MS/MS spectral matching in endo-metabolome (E,F) and exo-metabolome (G). (H,I) Chemical structure, relative abundance and co-elution of kynurenic acid (H) and *N*-acetyltyramine (I). Data are presented as mean ± SE. Unpaired *t* test was used to determine significance differences, \*\**p*<0.01, \*\*\**p*<0.001, \*\*\*\**p*<0.0001. **Sm**, *Skeletonema marinoi*; **Vc**, *Vibrio cyclitrophicus* HSW24; **CS1**, *Marinobacter adhaerens* CS1.

### *M. adhaerens* CS1 mitigates *V. cyclitrophicus* HSW24 induced growth inhibition and morphological damage in *S. marinoi*

Based on the pairwise interaction experiments, one significantly growth inhibiting bacterium *V. cyclitrophicus* and one significantly growth promoting bacterium *M. adhaerens* were selected for detailed study (Fig. 1A, B). In monoculture, *S. marinoi* exhibited growth from day 4 onwards, reached maximum cell counts at day 14 (RFU ∼3.2) and then declined by day 16. In coculture with *V. cyclitrophicus*, *S. marinoi* growth was significantly inhibited in comparison to the *S. marinoi* monoculture from day 6 onwards (*p*-value < 0.05). This inhibitory effect became more pronounced from day 8 (*p*-value < 0.001) onwards and reached a maximum on day 10 (RFU ∼1.0). On day 16, the fluorescence of *S. marinoi*-*V. cyclitrophicus* coculture treatment was reduced to a level comparable to that observed at beginning of the experiment (day 2) (Fig. 1A). In contrast, *M. adhaerens* had positive effect on *S. marinoi* growth. In coculture with *M. adhaerens*, *S. marinoi* growth was similar to that of monoculture during the early growth phase. But it increased from day 8 and became significantly higher (*p*-value < 0.001) than that of the monoculture from day 10 onwards (Fig. 1B). The maximum fluorescence was observed on day 14 (RFU ∼4.3).

To investigate whether *M. adhaerens* can modulate the inhibitory effect of *V. cyclitrophicus*, we set up a tripartite community consisting of *S. marinoi*, *V. cyclitrophicus* and *M. adhaerens*. Consistent with the pairwise coculture results, *V. cyclitrophicus* alone strongly inhibited the *S. marinoi* growth. When *M. adhaerens* was inoculated together with *V. cyclitrophicus* in equal cell number ratio (1:1; *M. adhaerens* : *V. cyclitrophicus*), *S. marinoi* growth was partially rescued (Fig. 1C). Increase in relative abundance of *M. adhaerens* further to ratios of 2:1, 3:1 (*M. adhaerens* : *V. cyclitrophicus)* supported *S. marinoi* growth in a ratio-dependent manner. All tripartite community treatments showed higher growth than the *S. marinoi*-*V. cyclitrophicus* coculture, indicating that *M. adhaerens* mitigates *V. cyclitrophicus* induced *S. marinoi* growth inhibition.

The microscopic observations qualitatively supported the fluorescence-based evaluation of the growth experiments (full time series of microscopic images shown in Supporting Fig. 3). At day 10, *S. marinoi* in monocultures appeared healthy with intact chains, uniform pigmentation and well-organized with relatively uniform cell morphology which reflects its optimal growth (Fig. 1D). In coculture with *V. cyclitrophicus*, *S. marinoi* exhibited altered morphology such as short chain, distorted cells, apparently cell lysis and numerous vesicle-like structures/cellular debris suggesting loss of cellular integrity (Fig. 1E). In contrast, coculture with *M. adhaerens* showed frequent chains, visibly higher cell abundance with uniform pigmentation which is consistent with positive effect of *M. adhaerens* on *S. marinoi* growth (Fig. 1F). In tripartite community, *S. marinoi* showed improved cell morphology, better cell dispersion, fewer lysed/damaged cells and fewer vesicle-like structures compared to *S. marinoi*-*V. cyclitrophicus* coculture (Fig. 1G). Most of the cells were present in small chains although few long chains were also observed.

**Figure 3.**
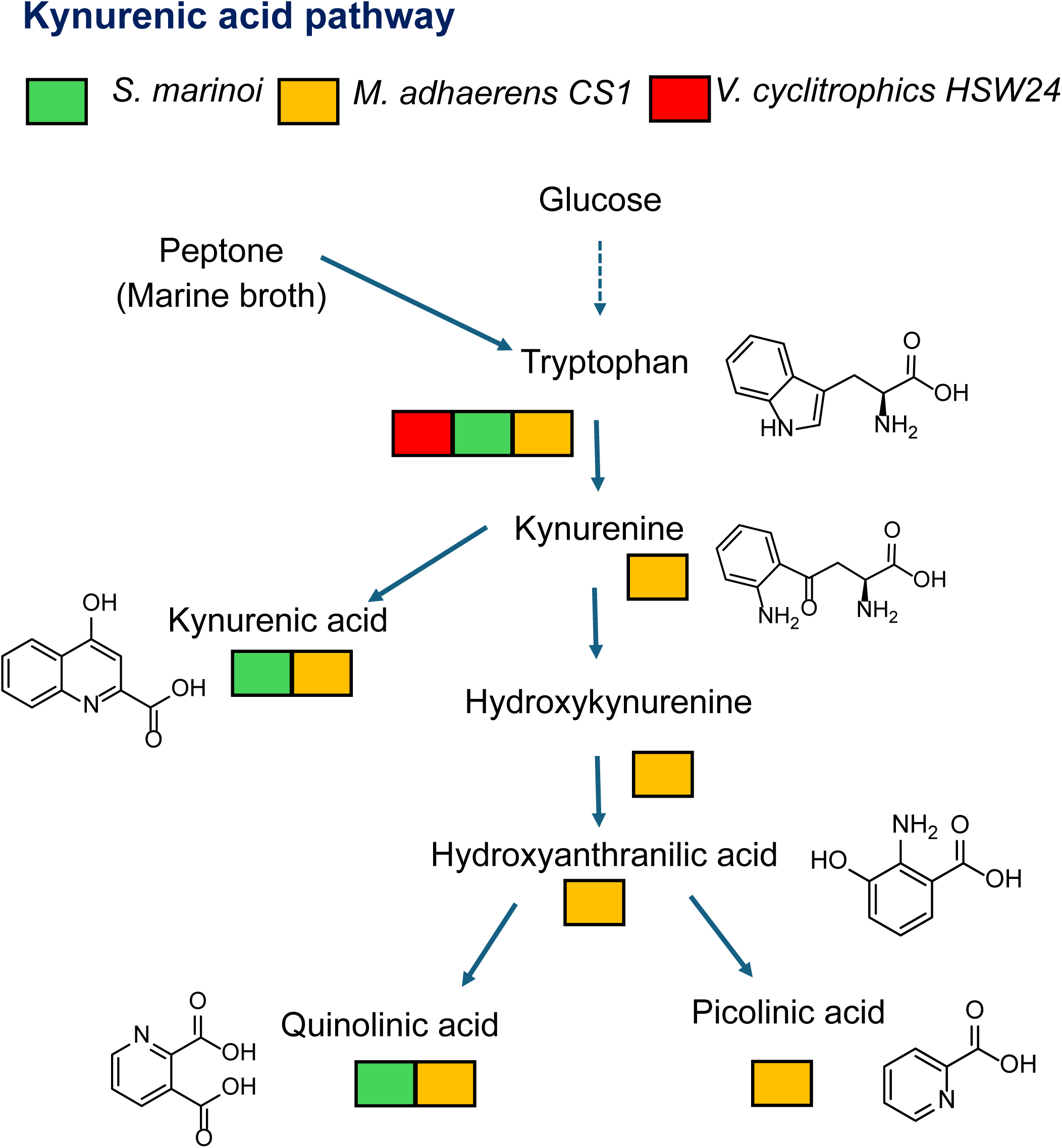
Kynurenic acid pathway-associated metabolites in the three interaction partners. Metabolite were detected in *S. marinoi* (green box), *M. adhaerens* CS1 (yellow box) and *V. cyclitrophicus* HSW24 (red box).

At day 16, *S. marinoi* monoculture retained uniform pigmentation but appeared mostly as isolated cells or in short chains (Fig. 1H). In contrast, very few *S. marinoi* cells with distorted morphology were observed in coculture with *V. cyclitrophicus* (Fig. 1I). Both *M. adhaerens* coculture and community showed visibly higher *S. marinoi* cells with healthier morphology compared to *S. marinoi* cocultured with *V. cyclitrophicus* (Fig. 1J, K). Growth and microcopy data suggest that *V. cyclitrophicus* inhibits *S. marinoi* growth and compromises cellular integrity whereas, *M. adhaerens* promotes growth and plays role in *S. marinoi* protection from *V. cyclitrophicus* mediated damage. Additionally, metabolomic changes associated with *S. marinoi - V. cyclitrophicus* coculture and bioassays with selected extracellular purine metabolites are provided in the supplementary data (Supporting Fig. 4,5).

**Figure 4:**
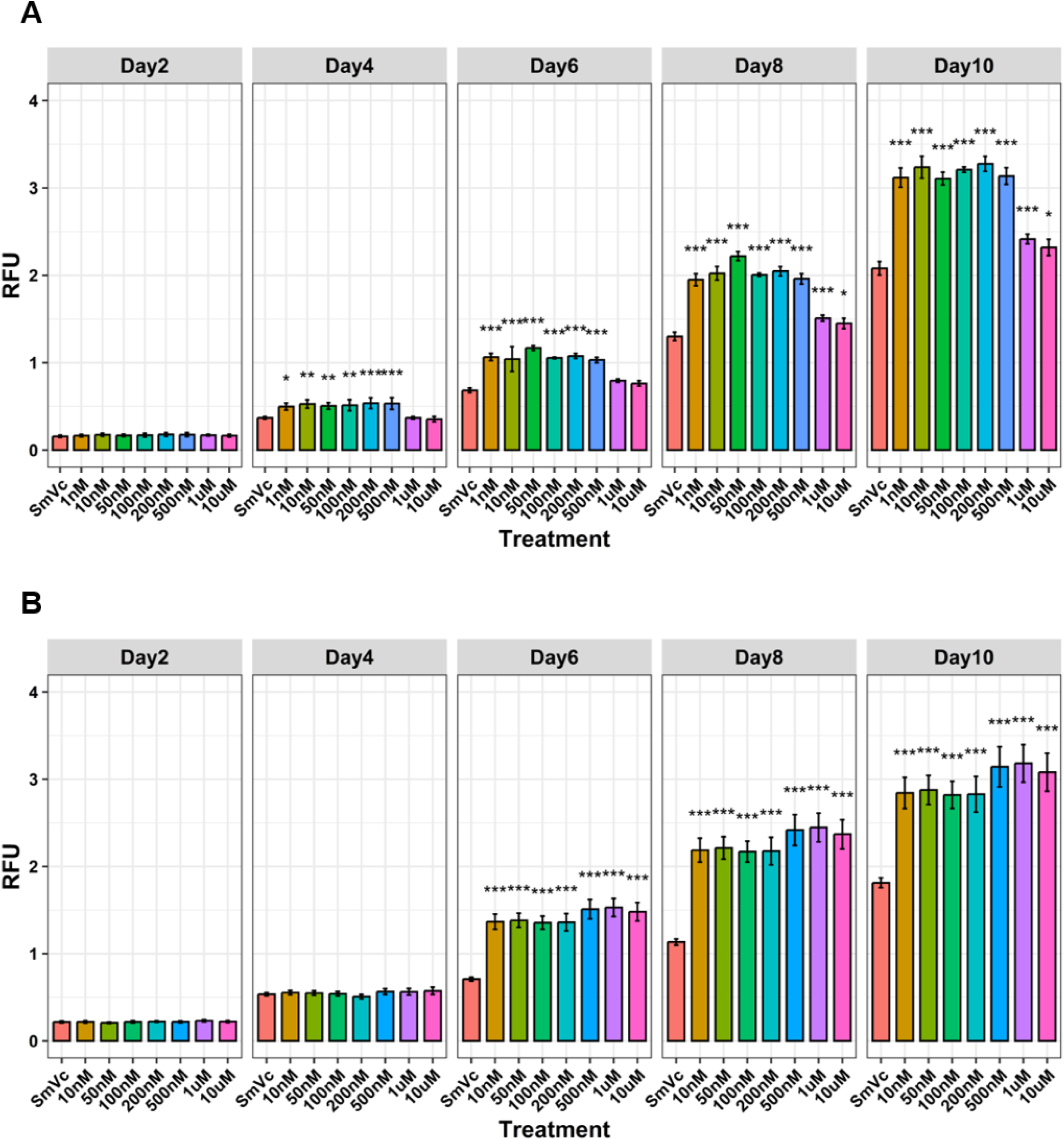
Effects of kynurenic acid and *N*-acetyltyramine supplementation on *S. marinoi* growth in coculture with *V. cyclitrophicus* HSW24. (A-B) Growth of *S. marinoi* in coculture with *V. cyclitrophicus* (SmVc) following supplementation with kynurenic acid at concentrations ranging from 1 nM to 10 µM (A) and *N*-acetyltyramine at concentrations ranging from 10 nM to 10 µM (B). Algal fluorescence was used as a proxy for *S. marinoi* growth and was measured on days 2, 4, 6, 8, and 10. Data are presented as mean ± SE from (n=8) biological replicates. Statistical significance was determined at each time point by ANOVA followed by Tukey multiple comparison test, * indicates *p*<0.05, ** indicates *p*<0.01, *** indicates *p*<0.001.

**Figure 5.**
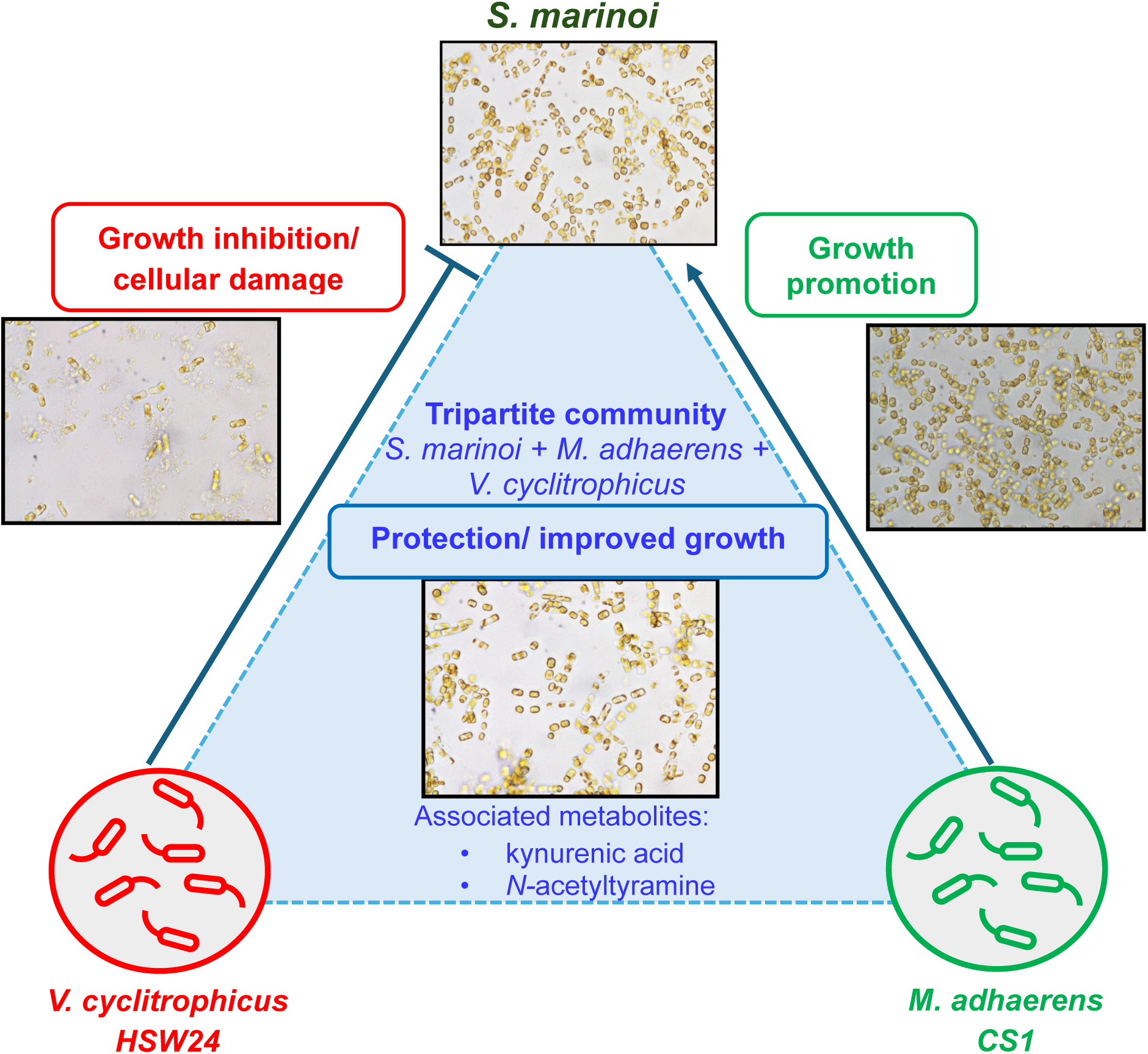
Conceptual model showing *S. marinoi* growth modulation by *M. adhaerens* in a tripartite community from *V. cyclitrophicus* HSW24. *V. cyclitrophicus* negatively affects *S. marino* growth and causes cellular damage in pairwise interaction whereas *M. adhaerens* protects *S. marinoi* growth in tripartite community. The model graph also highlights kynurenic acid and *N*-acetyltyramine as associated metabolites in protective community interaction.

### Metabolic changes associated with the protection of *S. marinoi* from the algicidal bacterium *V. cyclitrophicus* HSW24 in a tripartite community

We compared the endo- and exo-metabolome of *S. marinoi* grown in the tripartite community with *S. marinoi* grown in monoculture to identify candidate metabolites associated with the protection of *S. marinoi* from *V. cyclitrophicus*. The principal component analysis (PCA) score plot showed clear separation between the control and treatment groups on day 10. For the endo-metabolome data, PC1 and PC2 explained 48.3% and 15.2% of the total variance, respectively, whereas for the exo-metabolome data, they explained 69.7% and 9.8%, respectively (Fig. 2A,B).

The volcano plot analysis was performed to study differentially abundant metabolites between *S. marinoi* monoculture (control) and *S. marinoi* in a tripartite community (treatment) (Fig. 2C,D). In the endo-metabolome, 428 features were differentially regulated, out of which 264 features were significantly up-regulated and 164 features were significantly down-regulated in the tripartite community (Fig. 2C). These features were annotated based on elemental composition, *m*/*z*, adduct and retention time using the Compound Discoverer software. Further, the MS/MS spectral matching against mzCloud in Compound Discoverer provided putative annotations of features assigned as ethylbenzaldehyde and hydroxybenzaldehyde which showed significantly higher abundance in the tripartite community than in the *S. marinoi* monoculture (Fig. 2E,F). The exo-metabolome revealed a total of 379 differentially regulated features, out of which 336 metabolites were significantly up-regulated and 43 features were down-regulated (Fig. 2D). The MS/MS analysis in positive ionization mode revealed the up-regulation of quinolines such as hydroxyquinoline (Fig. 2G), methylquinoline and isoquinoline (Supporting Fig. 6A,B). Further, the up-regulation of kynurenic acid, a metabolite derived from tryptophan and *N*-acetyltyramine was observed. Both compounds were not up-regulated if bipartite cultures were compared to *S. marinoi* monocultures.The metabolites were further confirmed by matching MS/MS spectra and retention time with authentic standards (Fig. 4H,I and Supporting Fig. 7A,B).

### Detection of kynurenic acid pathway associated metabolites in *S. marinoi, V. cyclitrophicus and M. adherence*

Kynurenic acid is a product of the kynurenine pathway of tryptophan metabolism. Therefore, we screened the mass spectrometry datasets of *S. marinoi*, *V. cyclitrophicus* and *M. adhaerens* extracts for the features with accurate masses consistent with reported metabolites associated with this pathway (Fig. 3). The pathway starts from tryptophan which is converted into kynurenine, which is further converted to kynurenic acid. Kynurenine can also be converted into hydroxykynurenine and hydroxyanthranilic acid which is further catabolized into quinolinic acid or picolinic acid. We found that the precursor was present in all three organisms whereas kynurenic acid was detected only in *S. marinoi* and *M. adhaerens*. These findings suggest the presence of pathway-associated metabolites but do not demonstrate pathway activity or metabolic flux.

### Kynurenic acid and *N*-acetyltryamine protect *S. marinoi* from *V. cyclitrophicus*

We performed well plate assays to evaluate the effect of kynurenic acid and *N*-acetyltyramine on *S. marinoi* growth in the presence of *V. cyclitrophicus*. We tested a broad concentration range to cover the approximate concentrations determined by external calibration (Supporting Fig. 8A, B). The concentration of kynurenic acid and *N*-acetyltyramine were estimated to be 109 ± 6 nM and 441 ± 19 nM respectively in the exo-metabolome of the tripartite community. These concentrations were considered as semi-quantitative, since isotopically labelled internal standards were not prepared. To assess the protective function of kynurenic acid and *N*-acetyltyramine, the concentrations tested in the experiments were selected to extend below and above the concentration estimated.

For the well plate assay with kynurenic acid, we tested a range of concentrations from 1 nM to 10 µM and used fluorescence as a proxy to measure the *S. marinoi* growth up to day 10. On day 2, the growth for all treatments was low and comparable. The supplementation of kynurenic acid resulted in a significant increase in *S. marinoi* growth from day 4 onwards (Fig. 4A) compared to the *S. marinoi* - *V. cyclitrophicus*

(SmVc) coculture treatment. The tested concentration of 1 nM to 500 nM broadly showed plateau like response with no clear concentration dependent pattern. In contrast, the higher concentrations of 1 µM and 10 µM kynurenic acid did not show any effect till day 6. The growth increased significantly at day 8 and day 10 but the RFU values remain lower as compared to other tested concentrations.

For *N*-acetyltyramine, we tested a range of concentrations from 10 nM to 10 µM and this supplementation showed significantly higher RFU relative to *S. marinoi* - *V. cyclitrophicus* (SmVc) coculture over time. At day 2, the RFU values of all treatments were similar but from day 6 onwards *N*-acetyltyramine supplemented cultures showed a significant increase in growth (Fig. 4B). Like kynurenic acid, *N*-acetyltyramine led to a plateau-like effect with most of the tested concentrations. The well -plate assay with kynurenic acid and *N*-acetyltyramine clearly shows that they play a role in protection of *S. marinoi* from algicidal bacteria *V. cyclitrophicus* however, the mechanism underlying this protection remains unknown.

## DISCUSSION

Here we show that chemically mediated interactions shape multipartite interactions between bacteria and the diatom *S. marinoi*. The algicidal effect of *V. cyclitrophicus* can be alleviated by *M. adhaerens*, a bacterium that also supports *S. marinoi* growth in dipartite interactions. Using comparative metabolomics, we identify kynurenic acid and *N*-acetyltryamine from *M. adhaerens* that protect *S. marinoi* from *V. cyclitrophicus* in bioassays.

While the *S. marinoi* monoculture showed the expected growth pattern of a batch culture, co-culture with *V. cyclitrophicus* led to a significantly lower growth and earlier culture decline. Microscopic observations show that this process goes along with fragmentation of chains and lysis of *S. marinoi* cells from day 8. This loss of cellular integrity suggests that *V. cyclitrophicus* exerts an algicidal effect rather than merely acting by nutrient competition. To the best of our knowledge, algicidal activity of *V. cyclitrophicus* against a planktonic diatom has not been reported previously. This strain was isolated from seawater from Helgoland, where *S. marinoi* co-occurs, supporting the ecological plausibility of their interaction. Algicidal activity of members of the genus *Vibrio* is documented and *V. cyclitrophicus* that acts against a planktonic diatom extends the group of algicidal *Vibrio spp.* [15, 24]. This result is in consonance with growing evidence that antagonistic bacteria are a central component of diatom-associated microbiomes, particularly during periods of bloom aging or host stress, where algal exudation of metabolites shifts and chemical environment can become favorable for opportunistic antagonists [25, 26].

*M. adhaerens*, in contrast, supports growth of *S. marinoi*. Members of *Marinobacter* are common associates of marine algae. These bacteria can even be present in decade old *Skeletonema* cultures. They grow on and respond to algal metabolites, colonize the phycosphere and influence algal performance [27, 28]. The diatom growth modulating behaviour of members of the *Marinobacter* genus is well documented [29–32]. *M. adhaerens* inhibits the growth of the diatom *Coscinodiscus radiatus* whereas other *Marinobacter* species, such as *M. salarius* promote the growth of *S. marinoi* [2, 22]. In this study, not only support of *S. marinoi* growth, but also frequent diatom chain formation and uniform pigmentation were observed in the presence of *M. adhaerens*. Such beneficial processes can be mediated through nutrient remineralization, iron acquisition, vitamin supply, or detoxification of reactive compounds [2, 7].

In the tripartite synthetic community the bacterium *M. adhaerens* significantly protects *S. marinoi* from antagonistic *V. cyclitrophicus* although the protection did not fully restore the performance compared to the growth of the algae alone. The partial rescue of *S. marinoi* by *M. adhaerens* and the gradual increase in protection at higher *M. adhaerens* : *V. cyclitrophicus* ratios shows that the protection is abundance dependent. The algal microbiome function is thus not determined by pairwise interactions alone. Additional bacterial partners can change the outcome of antagonism by interfering with harmful bacteria, improving host stress tolerance, or reshaping the chemical environment around the host [7, 11, 33]. The recent finding that a mutualistic bacterium can rescue a green alga from bacterial antagonism in a tripartite culture further supports the ecological relevance of such protective effects [17].

Given the concentration-dependence of the rescue effect we reasoned that chemical factors might be involved. Therefore, we set out to identify candidate metabolites in a comparative metabolomics experiment. We searched for rescue molecules that would be more abundant in the tripartite community compared to the algae alone. Given the need for a metabolic exchange between the partners we focused on metabolites that are released into the water, i.e. members of the exo-metabolome. Indeed, the exo-metabolome of the tripartite community contained several metabolites that were not abundant in the *S. marinoi* metabolome or the metabolome of *the S. marinoi-V. cyclitrophicus* coculture (data not shown).

The exo-metabolome of the tripartite community was significantly enriched in kynurenic acid and *N*-acetyltyramine, with the identities of both metabolites unambiguously confirmed using synthetic standards. Kynurenic acid is a tryptophan-derived metabolite and its function in algae remains largely unexplored. Kynurenic acid can act as a molecule promoting mutualism in algae-bacteria interactions and it has been reported that bacterial derived kynurenic acid can protect algae from cadmium stress [36]. In the present experiment, exogenously administered kynurenic acid significantly supported *S. marinoi* growth in the presence of *V. cyclitrophicus*, indicating that its protective effect is not restricted to abiotic stress but also to interactions with antagonistic bacteria. The assay showed a pronounced rescuing effect of kynurenic acid at lower concentrations with weaker responses at the highest doses, suggesting a non-linear dose dependence which is consistent with the behaviour of small signalling or protective metabolites that become toxic at elevated concentrations. It has to be noted that the concentration within the phycosphere might differ from that determined here in the dissolved fraction of the exo-metabolome [3]. Therefore, the concentration estimated from the extract should be interpreted as an average value and may not necessarily reflect the local concentrations experienced by cells within the phycosphere. Therefore, we tested a broad concentration range that is expected to cover the actual effective concentrations. Working with a tripartite community it cannot be safely stated which partner contributes the respective metabolites detected in the exo-metabolome. Pathway analysis however revealed that upstream and downstream metabolites related to kynurenic acid biosynthesis are found in *M. adhaerens*, while they were not detected in both *S. marinoi* and *V. cyclitrophicus*, making it likely that *M. adhaerens* is the producer of the metabolite.

Another molecule that was significantly up-regulated in the exo-metabolome of the tripartite community is *N*-acetyltyramine. This metabolite has reported quorum-sensing inhibitory and anti-virulence functions and is thus also a promising candidate for the mediation of protective effects in the tripartite community [37]. The bioassay with *N*-acetyltyramine showed a significant effect from day 6 onward. Even low nanomolar concentrations were sufficient for partial restauration of the *S. marinoi* growth in the presence of *V. cyclitrophicus*, which supports the idea that *M. adhaerens* protection may involve suppression of *V. cyclitrophicus* communication-dependent antagonistic traits. Together, these results suggest that *M. adhaerens* protects *S. marinoi* by providing several metabolites including kynurenic acid and *N*-acetyltyramine that may enhance *S. marinoi* tolerance or reduce *V. cyclitrophicus* virulence. Thus, protection in the tripartite community appears to arise from distinctive metabolic changes that cannot be predicted from pairwise interactions alone. This finding is consistent with previous studies demonstrating that the outcome of algal–bacterial interactions is strongly influenced by community composition, with additional microbial partners capable of altering the net effects of harmful bacteria on the host [17, 33]. More broadly, it supports the view that plankton microbiomes operate through complex networks of metabolite exchange and higher-order interactions rather than through isolated pairwise relationships[38].

In addition, quinolines were significantly up-regulated in the exo-metabolome of the tripartite culture. These metabolites have metal-chelating, redox-activity and antimicrobial properties making them plausible candidates for a regulatory activity in the tripartite community [34]. Although the function of these quinoline related molecules remain unresolved in the present study, their accumulation suggests a potential role in chemically mediated interactions that has to be subject to further bioassays.

## CONCLUSION

Our findings demonstrate that algal–bacterial interactions cannot be understood from pairwise interactions alone. While *V. cyclitrophicus* strongly inhibited *S. marinoi* growth and altered cell morphology, the addition of *M. adhaerens* mitigated these effects in an abundance-dependent manner (Fig. 5). Metabolomic analyses revealed concomitant remodelling of endo- and exometabolite profiles, with kynurenic acid and *N*-acetyltyramine emerging as candidate mediators of protection. Exogenous addition of both compounds alleviated the antagonistic effect of *V. cyclitrophicus*. Together, these results suggest that algal-associated bacteria can provide a protective function beyond nutritional support, highlighting the importance of community context in shaping diatom resilience to bacterial antagonism.

## ACKNOWLEDGEMENT

We thank Gabriel Jahn for his support in this research during his Bachelor’s Thesis.

## SUPPLEMENTARY DATA

Supplementary data is available online only

Supplementary file 1

## AUTHOR CONTRIBUTIONS

S.A.S. and G.P. conceptualized the project. S.A.S., C.Z., V.N., R.Y. and G.P. developed methodology. S.A.S. conducted experimental investigations. C.Z. contributed to metabolomics experiments and analytical support. S.A.S., C.Z., V.N. and R.Y. conducted formal analysis and interpretation. S.A.S and G.,P. wrote the original manuscript draft. G.P. supervised the project and acquired funding.

## DATA AVAILABILITY

All supporting data related to the present study are provided within the manuscript, in the supplementary files or deposited in public repositories. The 16S rRNA gene sequence of isolated bacteria in the present study has been submitted to GenBank with accession number PZ671326. Metabolomics data generated in the present study are available via MetaboLights with identifier MTBLS14988.

