## Supplementary data and figures for "Algae–bacteria associations provide metabolite-mediated protection against algicidal bacteria in a tripartite plankton community"

Short title: protective tripartite microbial interactions

Shahrukh A. Siddiqui[1,2], Christian Zerfaß[1], Vera Nikitashina[1,2], Ruyi Yu[1], Georg Pohnert[1,2*].

[1]Department of Bioorganic Analytics, Institute for Inorganic and Analytical Chemistry, Friedrich Schiller University, Jena, Germany.

[2]Cluster of Excellence Balance of the Microverse, Friedrich Schiller University, Jena, Germany.

**Supporting Table 1.** Bacterial isolates used in the present study.

| S. No | Strain | Species | Accession Number (NCBI) |
|---|---|---|---|
| 01 | DG893 | *Marinobacter algicola* | AY258110.2 |
| 02 | CR1 | *Pseudoalteromonas sp.* | MZ605002 |
| 03 | Rose1 | *Roseovarious sp.* | MZ605004 |
| 04 | CIP105210 | *Phaeobacter gallaeciensis* | KC176239.1 |
| 05 | DSM17395 | *Phaeobacter inhibens* | KC176241.1 |
| 06 | Marino1 | *Marinobacter sp.* | MZ605006 |
| 07 | Cro1 | *Croceibacter Sp.* | MZ605003 |
| 08 | Mari1 | *Maribacter sp.* | MZ605005 |
| 09 | CS1 | *Marinobacter adhaerens* | MZ605007 |
| 10 | HSW24 | *Vibrio cyclitrophicus*-HSW24 | PZ671326 |

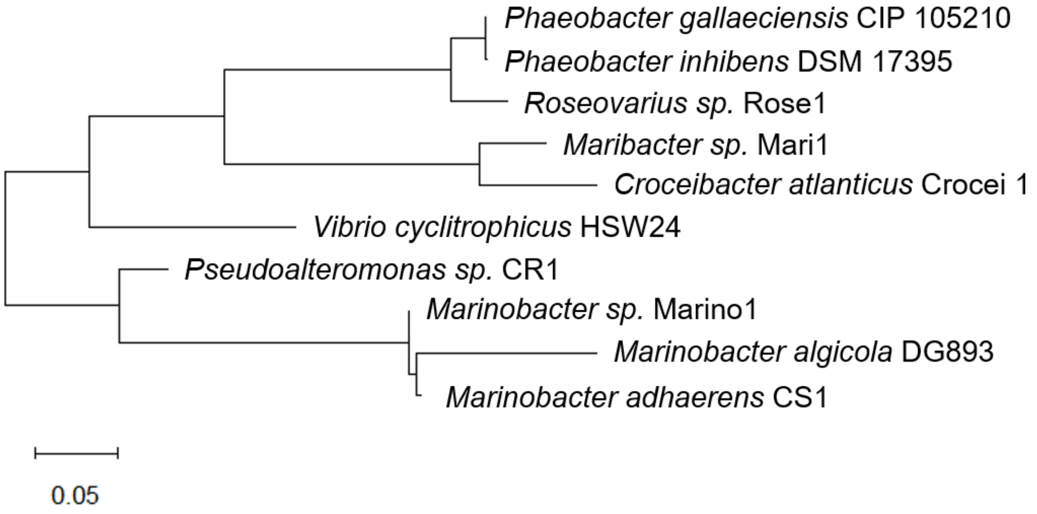


**Supporting Figure 1.** 16S rRNA gene-based phylogenetic tree of the bacterial isolates used in this study. The tree illustrates the phylogenetic relationships among the screened marine bacterial strains. The scale bar indicates 0.05 substitutions per nucleotide position.

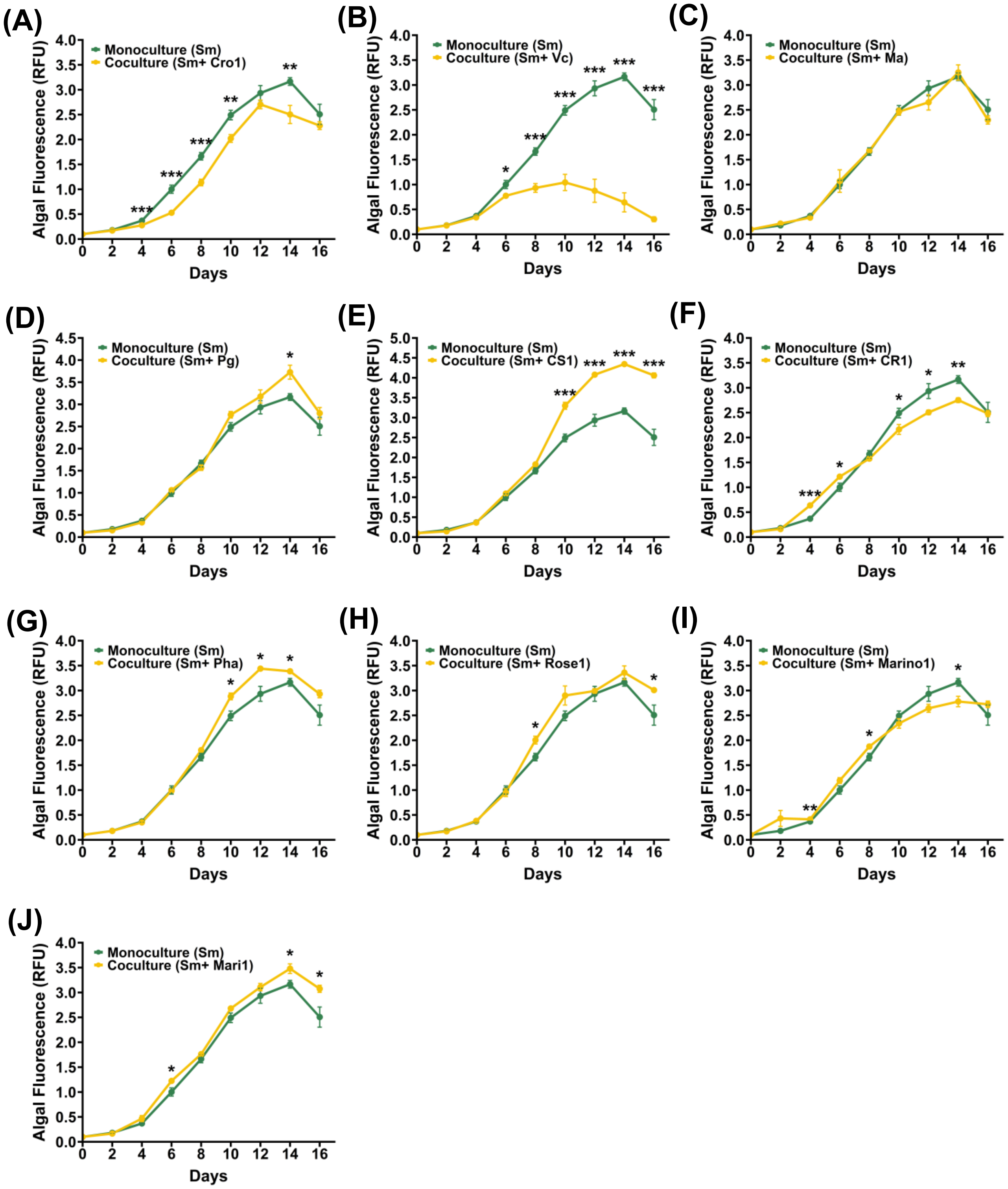


**Supporting Figure 2.** *S. marinoi* (Sm) pair-wise interaction with bacteria for screening test. *S. marinoi* growth were observed for 16 days either in monoculture (Sm, green line) or in coculture with individual bacterial strains (yellow line). The chlorophyll fluorescence (RFU) was used as proxy to growth. Significant differences between monoculture and coculture at individual time points were assessed using student's *t*-test and are indicated by asterisks ($P < 0.05$, *$P < 0.01$, **$P < 0.001$). Data are presented as mean ± SD of 5 biological replicates. **Sm**, *Skeletonema marinoi*; Cro1, *Croceibacter Sp*, **Vc**, *Vibrio cyclitrophicus* HSW24; **Ma,** *Marinobacter algicola*; **Pg**, *Phaeobacter gallaeciensis;* **CS1**, *Marinobacter*

*adhaerens*; **CR1**, *Pseoalteromonas sp*.; **Pha**, Phaeobacter inhibens*;* **Rose1***,* Roseovariouss sp.*;* **Marino 1***,* Marinobacter sp,*;* **Mari1,** *Maribacter sp.*

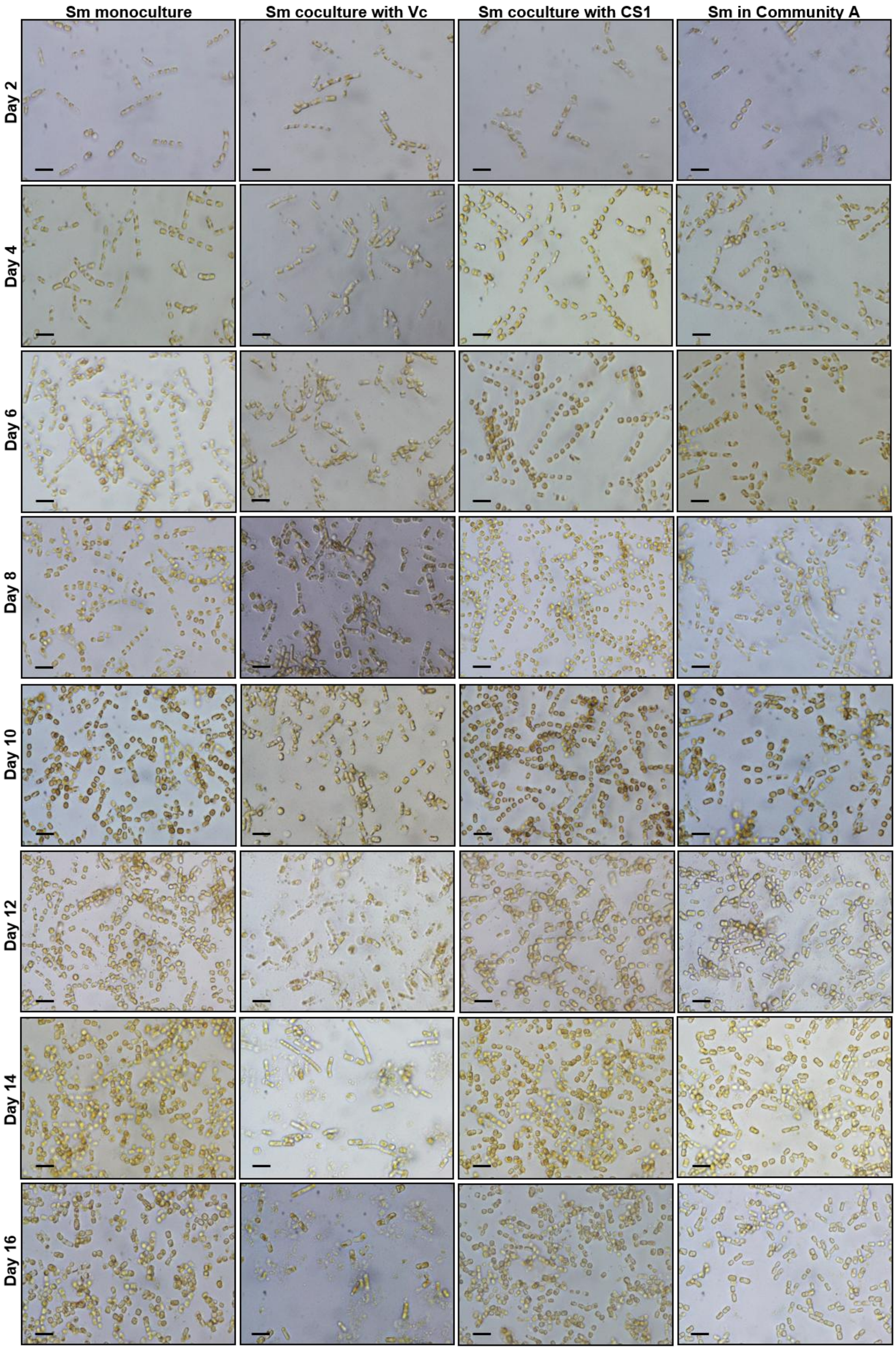

**Supporting Figure 3.** Microscopy images of *S. marinoi* (Sm) monoculture (column 1), coculture with Vc (column 2), coculture with CS1 (column 3) and community A (column4). Scale bar represents 20µm.

### Metabolomic changes associated with *S. marinoi* - *V. cyclitrophicus* HSW24 coculture

To investigate the metabolic response of *V. cyclitrophicus* on *S. marinoi*, we compared the endo and exo-metabolome of *S. marinoi* grown alone (control) and in coculture with *V. cyclitrophicus* (SmVc; treatment) on day 10. Principal Component Analysis (PCA) score plot showed clear separation between control and treatment groups. For endo-metabolome, *S. marinoi* and *S. marinoi* cocultured with *V. cyclitrophicus* formed two distinct clusters in the PCA score plot with PC1 and PC2 explaining 40% and 16.5% of the total variance respectively (Supporting Fig. 4A). *S. marinoi* replicates clustered tightly whereas, *S. marinoi* cocultures with *V. cyclitrophicus* replicates were dispersed indicating enhanced variability in coculture induced by Vc. Similarly, for exo-metabolome, *S. marinoi* and *S. marinoi* cocultured with *V. cyclitrophicus* also formed two distinct clusters in PCA score plot and exhibited 57.9% and 21.1% of the total variance in PC1 and PC2 respectively (Supporting Fig. 4B).

The volcano plot analysis was performed to investigate the differentially regulated metabolites between *S. marinoi* and *S. marinoi* cocultured with *V. cyclitrophicus* in the endo- and exo-metabolome (Supporting Fig. 4C, D). The metabolites with fold change value of greater than 2 or less than 0.5 in a volcano plot at FDR adjusted $p$-< 0.05 were considered differentially regulated. In the endo-metabolome, a total of 202 features were differentially regulated (Supporting Fig. 4C), out of which 136 features were significantly up-regulated and 66 features were significantly down-regulated. The MS/MS spectra analysis using SIRIUS in positive mode provided the putative annotations for several features associated with tetrapyrrole and electron transport chain, such as herderoporphyrin, pheophorbide, a tetrapyrole derivative and ubiquinone (Supporting Fig. 4E-H).

In the exo-metabolome, a total of 338 features were differentially regulated, with 67 significantly up-regulated features and 271 significantly down-regulated features. The MS/MS spectral matching against mzCloud library in compound discoverer revealed the putative annotation of up-regulated features including adenine, adenosine, deoxy-adenosine and methyl-indoleacetate (Supporting Fig. 4I-L). Further, adenine and 2-

deoxyadenosine were confirmed with authentic standards, whereas methyl indoleacetate remain putatively annotated.

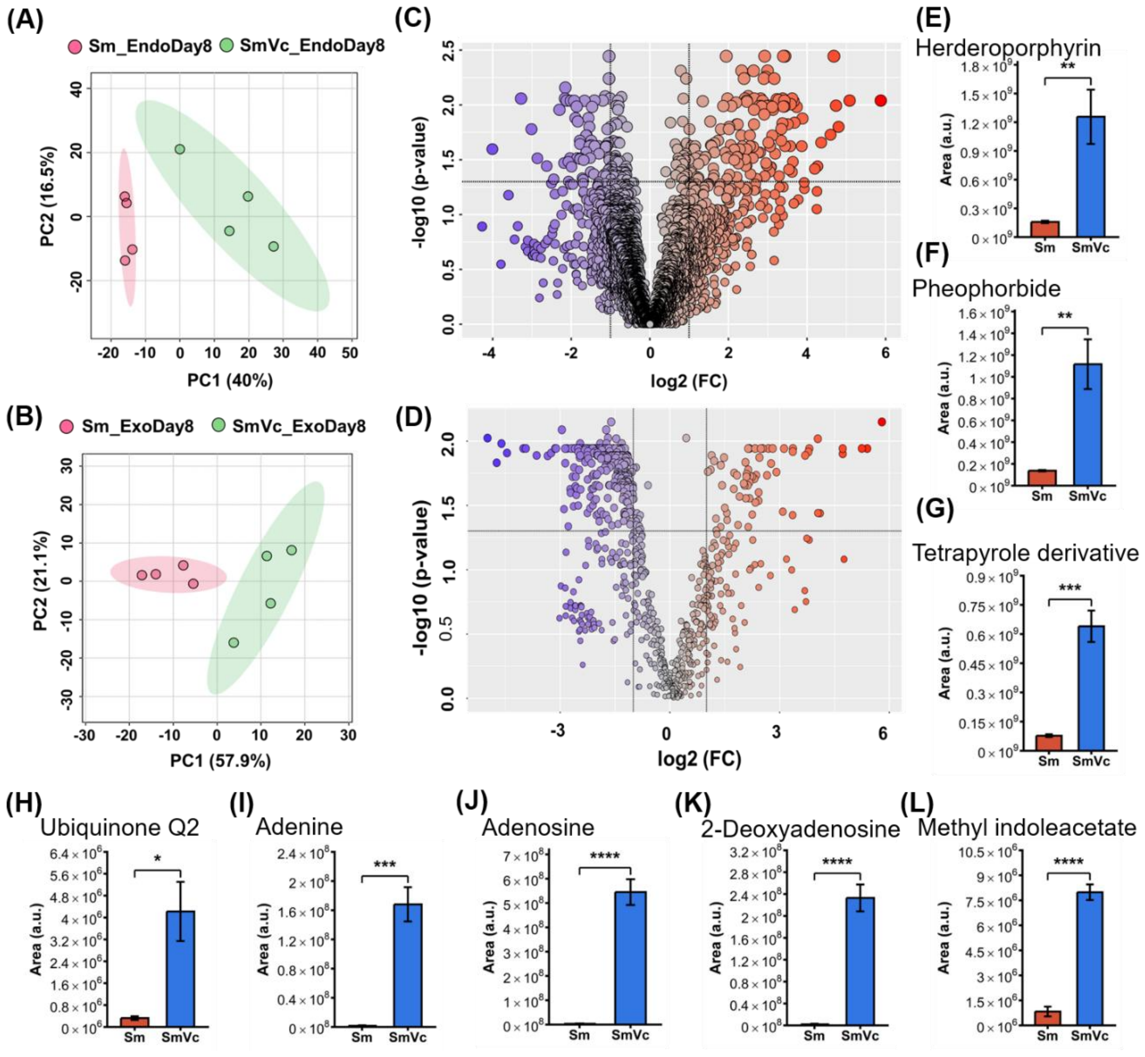


**Supporting Figure 4.** Metabolomic changes in *S. marinoi* cocultured with *V. cyclitrophicus*. (a,b) Principal Component Analysis of endo-metabolome (a), exo-metabolome (b) of *S. marinoi* (Sm; control) grown alone or in co-culture with *V. cyclitrophicus* (SmVc; treatment). (c,d) Volcano plots showing differentially abundant features in the endo-metabolome (c) and exo-metabolome (d). Red dots indicate up-regulation and blue dots indicate down-regulation. (e-h) Relative abundance of selected endo-metabolome features putatively annotated using MS/MS spectral data with SIRIUS. (i-j) Relative abundance of selected exo-metabolome metabolites annotated by MS/MS spectral matching against mzCloud library. Adenine and 2-deoxyadenosine were further confirmed by running authentic standards. Data are presented as mean ± SE. Unpaired *t* test was used to determine significance differences, * indicates $p<0.05$, ** $p<0.01$, *** $p<0.001$, **** $p<0.0001$.

**Effect of extracellular purine metabolites on *S. marinoi* growth**

Metabolomic analysis revealed the up-regulation of several purine nucleobases and nucleosides in the exo-metabolome of *S. marinoi* cocultured with *V. cyclitrophicus* (treatment) compared with *S. marinoi* monoculture (control). To test whether these compounds could directly affect *S. marinoi* growth. We selected adenine and 2-deoxyadenosine for supplementation bioassays because both were up-regulation in the *S. marinoi* cocultured with *V. cyclitrophicus* exo-metabolome and their identities were confirmed using authentic standards. Each compound was tested at 0.1 μM, 1 μM, 10 μM and 100 μM concentration on *S. marinoi* and growth was monitored for 10 days. Neither adenine nor 2-deoxyadenosine caused a significant change in *S. marinoi* growth at any of the tested concentrations (Supporting Fig. 5A,B).

(A)

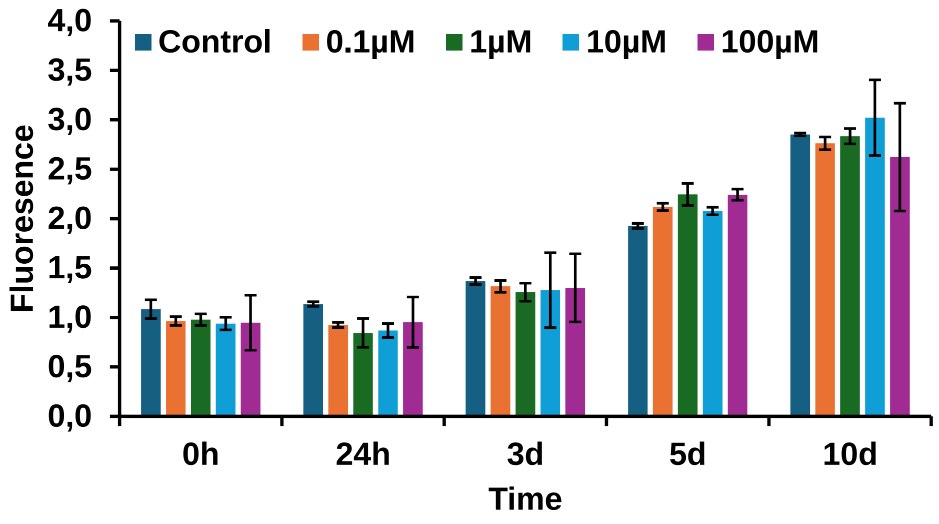


(B)

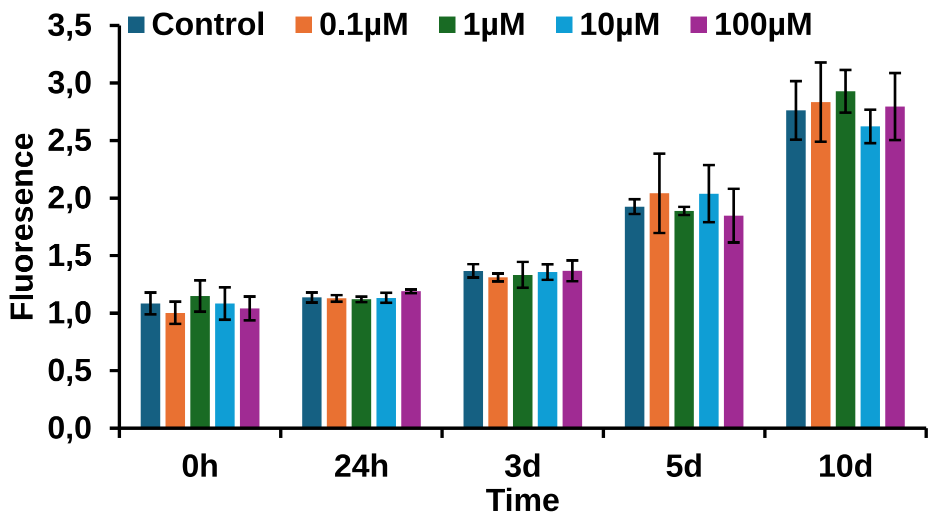

**Supporting Figure 5.** Effect of extracellular purine metabolites on *S. marinoi* (Sm) growth. (a) Growth of Sm after supplementation with adenine and (b) 2-deoxyadenosine at 0.1, 1, 10, and 100 μM. These compounds were tested because they showed higher relative abundance in the extracellular metabolome of SmVc compared with Sm monoculture and were confirmed using authentic standards. Growth was monitored for 10 days using chlorophyll a fluorescence as a proxy for algal growth. Data are presented as mean ± SE from biological replicates.

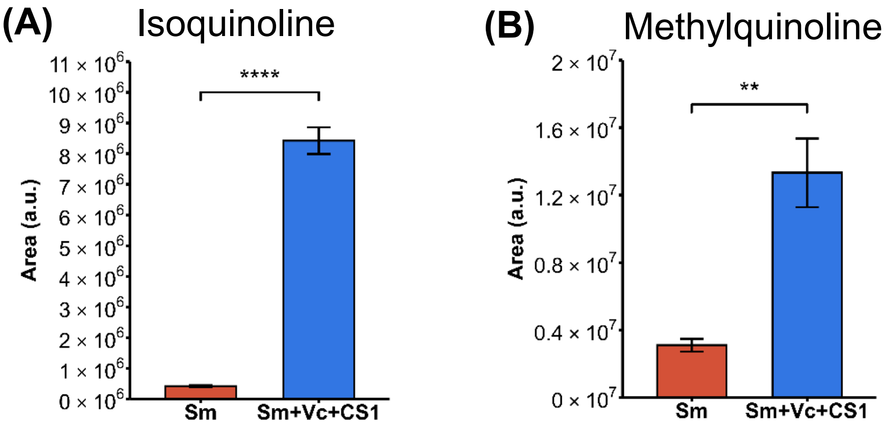


**Supporting Figure 6.** Relative peak area of (a) isoquinoline and (b) methylquinoline in exo-metabolome of tripartite community interaction. Data are presented as mean ± SE. **p<0.01, ***p<0.001, ****p<0.0001.

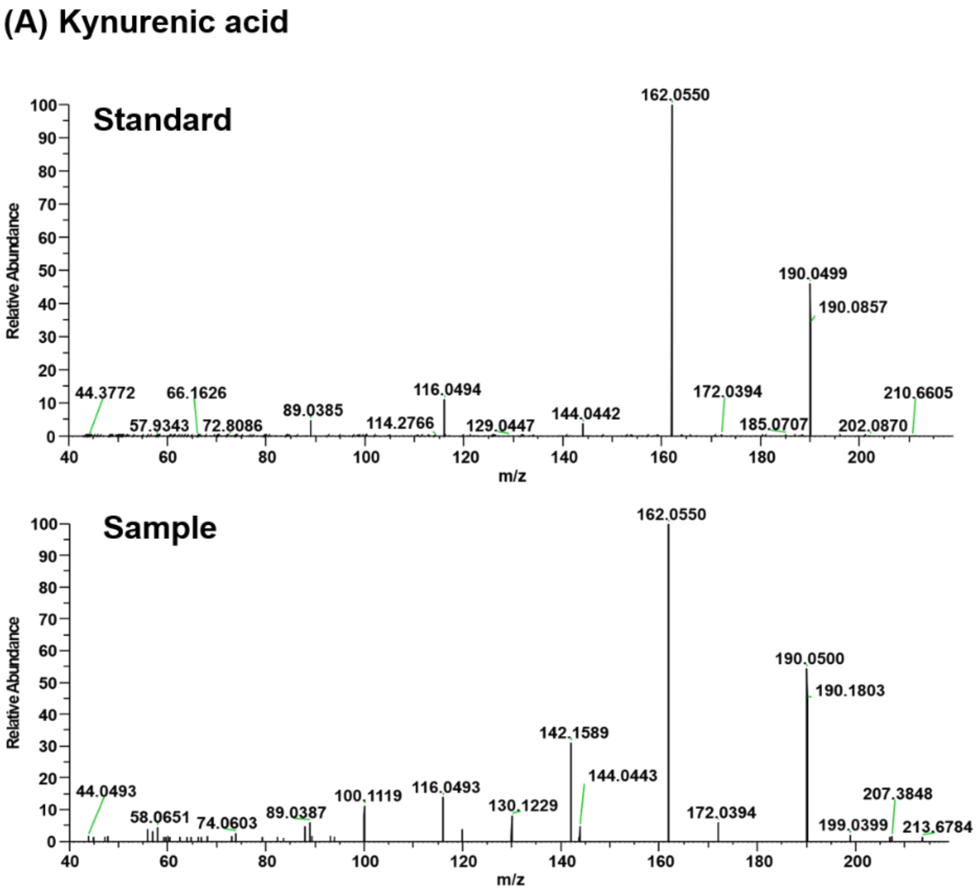

### (B) N-acetyltyramine

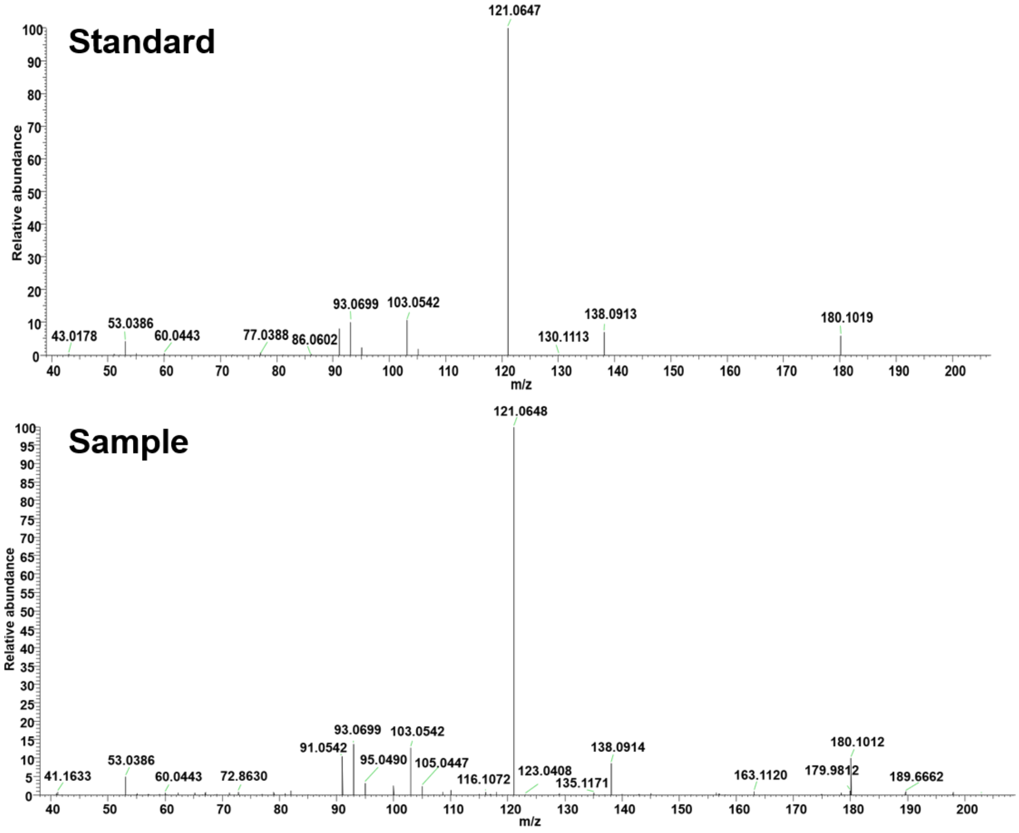


**Supporting Figure 7.** MS/MS spectral confirmation of kynurenic acid ans N-acetyltyramine. MS/MS spectra of authentic standards and corresponding sample features for (a) kynurenic acid and (b) N-acetyltyramine.

### (A) Kynurenic acid

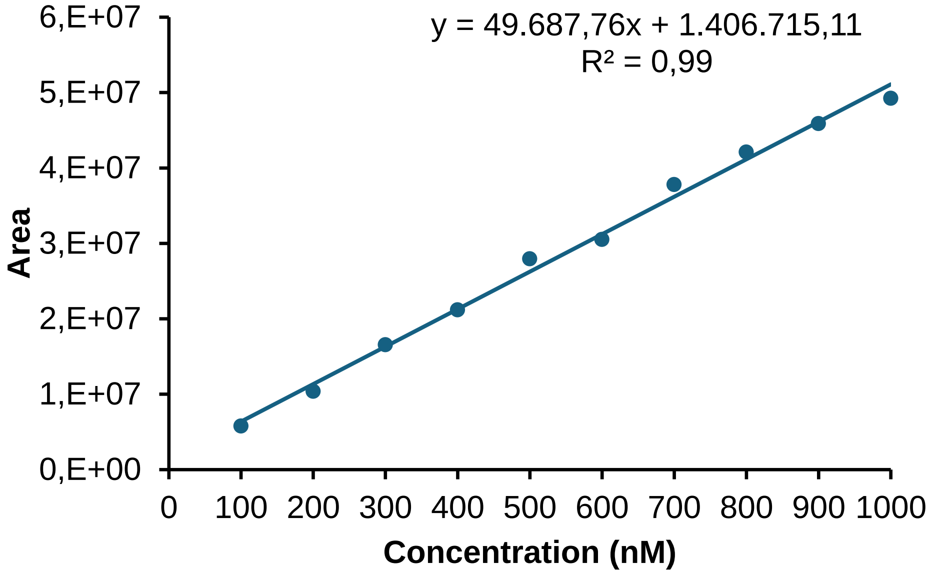

**(B)** N-acetyltyramine

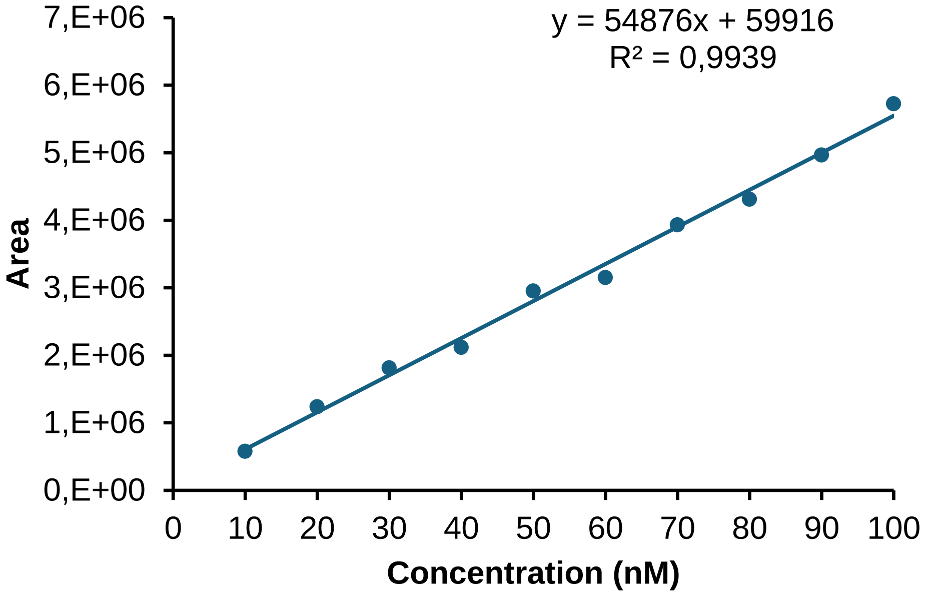


**Supporting Figure 8.** External calibration curve of (a) kynurenic acid and (b) N-acetyltyramine.
